# Psilocybin as a Treatment for Repetitive Mild Head Injury: Evidence from Neuroradiology and Molecular Biology

**DOI:** 10.1101/2025.02.03.636248

**Authors:** Eric K. Brengel, Bryce Axe, Ashwath Maheswari, Muhammad I. Abeer, Richard J. Ortiz, Taylor J. Woodward, Reagan Walhof, Rachel Utama, Courtney Sawada, Shreyas Balaji, Praveen P. Kulkarni, Heather B. Bradshaw, Michael A. Gitcho, Craig F. Ferris

## Abstract

Repetitive mild head injuries incurred while playing organized sports, during car accidents and falls, or in active military service are a major health problem. These head injuries induce cognitive, motor, and behavioral deficits that can last for months and even years with an increased risk of dementia, Parkinson’s disease, and chronic traumatic encephalopathy. There is no approved medical treatment for these types of head injuries. To this end, we tested the healing effects of the psychedelic psilocybin, as it is known to reduce neuroinflammation and enhance neuroplasticity. Using a model of mild repetitive head injury in adult female rats, we provide unprecedented data that psilocybin can reduce vasogenic edema, restore normal vascular reactivity and functional connectivity, reduce phosphorylated tau buildup, enhance levels of brain-derived neurotrophic factor and its receptor TrkB, and modulate lipid signaling molecules.

## Introduction

The Centers for Disease Control and Prevention report around 2.9 million people in the United States suffer from traumatic brain injury (TBI) every year, with 70-90% of these categorized as mild TBI^1, 2, 3^. In 2016, the total yearly healthcare expenses for nonfatal TBI exceeded $40.6 billion. This included $10.1 billion from private insurance, $22.5 billion from Medicare, and $8 billion from Medicaid^3^. There is an expanding literature on the behavioral and neurobiological consequences of mild head injuries that are incurred while playing organized sports, during car accidents and falls, or in active military service. Concussion following a single incident is difficult to detect and any associated cognitive and behavioral problems can resolve within hours of insult^4, 5^.

However, a more pernicious, long-lasting condition arises when the brain is exposed to repetitive mild traumatic brain injury (rmTBI)^6, 7^. Repetitive head impacts and rmTBI induce cognitive, motor, and behavioral deficits, which are more severe and protracted, and can last for months and even years^8, 9^ with an increased risk of dementia, Parkinson’s disease^10–14^, and chronic traumatic encephalopathy (CTE)^15, 16^. There are no approved treatments for repetitive head impacts, TBI, or rmTBI.

It has been suggested that the serotonergic hallucinogen psilocybin (PSI) could be used to treat brain injury given its known anti-inflammatory effects and its action as a promoter of neuroplasticity and cell growth^17^. A wide range of psychedelics, including PSI, are being evaluated for their potential therapeutic use in various psychiatric disorders^18^ including substance abuse^19^, severe depression^20, 21^, and anxiety^22^. To date, there are no reports of PSI being used to treat any type of head injury. In a recent functional MRI study, we evaluated the dose-dependent effects of PSI on brain activity in fully awake rats. From these studies we determined a 3.0 mg/kg dose of PSI was most effective in stimulating positive blood-oxygen-level-dependent (BOLD) changes in brain activity. In this study, we tested the efficacy of this dose of PSI in a closed-skull momentum exchange model of rmTBI wherein rats were impacted once each day for three consecutive days^23–27^. To make the model more relevant to the human experience, rats were impacted while fully awake and during the dark phase of the light-dark cycle, when they are normally active. There are no mortalities with this model. Absent are any contusions or damage to the skull. Instead, the only gross radiological evidence of head injury is swelling of the tissue above the skull at the impact site, i.e. a “bump on the head.” A single dose of PSI after each head impact significantly reduced the neuroradiological and molecular measures associated with head injury.

## Methods and Materials

### Animals

Adult female *(N =* 24) Wistar rats were purchased from Charles River Laboratories (Wilmington, MA, USA). Animals were housed in Plexiglas cages (two per cage) and maintained in ambient temperature (22–24°C). Animals were maintained on a reverse light-dark cycle with lights off at 09:00 and studied during the dark phase when they are normally active. All experiments were conducted between 10:00 and 18:00 to avoid the transitions between the light-dark cycles. Food and water were provided *ad libitum*. Rats were ca. nine months of age when head-impacted and imaged. All animals were acquired and cared for in accordance with the guidelines published in the NIH Guide for the Care and Use of Laboratory Animals. All methods and procedures described below were pre-approved by the Northeastern University Institutional Animal Care and Use Committee under protocol numbers 23-0407R and 24-0517R. Northeastern University’s animal care and use program and housing facilities are fully accredited by AAALAC International. The protocols used in this study followed the ARRIVE guidelines for reporting *in vivo* experiments in animal research^28^. Animals were monitored daily over the duration of the study for general health, including food and water consumption and body weight. A 15% loss in body weight was set as a humane endpoint. Female rats were divided into three experimental groups (*n* = 8; determined by *a priori* power analysis): 1) healthy sham controls injected with saline vehicle but given no head impact (SHAM-VEH), 2) head-impacted and injected with saline vehicle (rmTBI-VEH), and 3) head-impacted and injected with psilocybin (rmTBI-PSI).

### Overview

The study began with three consecutive days of mild head injury, psilocybin treatment, and head-twitch response observation. On day three, within an hour of head injury and treatment, blood plasma samples were collected for lipidomic analysis of peripheral biomarkers of mild head injury. This was immediately followed by the first magnetic resonance imaging (MRI) session. Cognitive and motor behaviors were tested on days 4-10. Acclimation for awake neuroimaging occurred on days 15-19 leading up to the second MRI session on day 22. The following day, brain tissue was collected for proteomic analysis. See **Fig 1a** for a timeline of experimental procedures.

**Fig 1.**
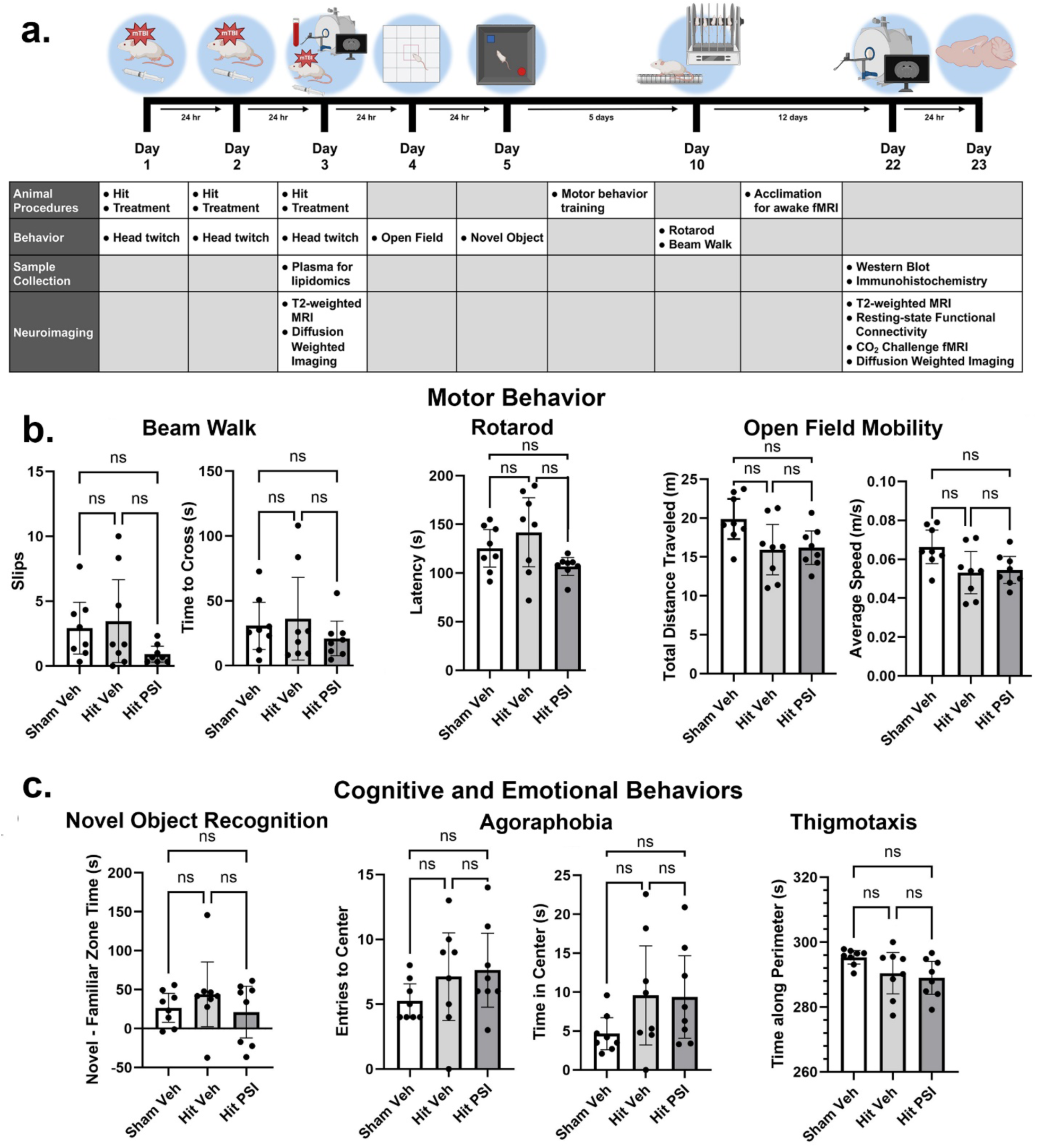
Experimental Protocol and Behavioral Assessment. **a.**) Timeline of experimental procedures. **b.**) Motor behavior on the Beam Walk, Rotarod, and Open Field tests. **c.**) Cognitive behavior from Novel Object Recognition and emotional behaviors agoraphobia and thigmotaxis collected from the Open Field test. All behavioral data were collected within one week of the last head impact. One-way ANOVAs showed no significant (ns) differences between any of the experimental groups for any assay.

### Repetitive Mild Head Injury

The momentum exchange model of mild head injury was developed by Viano et al.^29^ and further refined by Mychasiuk et al.^30^ and Hightower et al.^31^ to simulate the dynamics of sport-related concussion in the preclinical setting with fully conscious rodents. Each rat underwent this procedure once per day for three consecutive days. Prior to impact on the first day, all rats were treated with 0.1 mg/kg extended-release buprenorphine analgesic via subcutaneous injection to minimize pain for the duration of the three-day repetitive injury period. Rats were lightly anesthetized via 1-2% isoflurane inhalation to allow for setup in the momentum exchange apparatus. Under anesthesia, rats were secured with a bite bar and strapped to a wheeled cradle sitting atop a chassis to allow for linear and rotational acceleration upon impact. Once fully awake (typically within 1-2 minutes), a pneumatic pressure system is used to reliably propel a 50 g impactor toward the head at 7.4 m/s for a kinetic energy input of 1.37 J, simulating the rapid change in head velocity that occurs during concussions in the National Football League^29, 30^. With the head angled downward into the impact plane, all injuries were directed to the approximate area of Bregma. All rats demonstrated normal ambulatory behavior within seconds of being returned to the home cage, and no mortalities, seizures, loss of consciousness, skull fractures, contusions, or other complications were observed. SHAM-VEH control rats underwent all of the above procedures with the exception of head impact; when fully awake, these rats were removed from the apparatus and the impactor was not launched.

### Psilocybin administration and Head Twitch Response

Psilocybin was acquired through the National Institute on Drug Abuse (NIDA) and distributed by the Research Triangle Institute. PSI was prepared in sterile saline (0.9% NaCl) at a 3.0 mg/mL concentration for a dose of 3.0 mg/kg via intraperitoneal injection.

Rats were treated within 30 minutes of each head injury (once per day for three days). After each injection, rats were returned to the home cage and recorded from above for 10 minutes for quantification of the head twitch response (HTR), a behavioral indicator of psychoactive dosing. HTR and all subsequent behavioral assays were analyzed without outliers using one-way ANOVA in GraphPad Prism 10.0 software.

### Open Field

On day 4, within 24 hours of the last head impact, rats were tested in the Open Field under dim red-light illumination. A detailed description of the Open Field test in rats appears in previous publications^32, 33^. Animals were placed in a lidless black box (60.9 x 69.2 x 70.5 cm) and allowed to explore for 5 minutes while recorded from above.

Recordings were processed and data were measured using ANY-maze 7.00 software. In processing, the arena was divided into a peripheral zone (18 cm from the walls) and a central zone (20% of the arena). Agoraphobia (time spent in the center and number of entries to the center), thigmotaxis (time spent along the perimeter), average speed, and total distance traveled were measured and compared.

### Novel Object Recognition

On day 5, the Novel Object Recognition task was used to evaluate episodic learning and memory^34–37^. Testing was conducted in the same lidless black box under dim red-light illumination, to which rats were acclimated during Open Field testing the previous day. Rats were first given a 5-minute habituation phase inside the empty box. Rats were then given a 5-minute familiarization phase to explore the box with two identical objects placed in diagonal corners. Lastly, rats returned for the 5-minute test phase, where one familiar object and one new object were presented in the same positions as the familiarization phase. Rats were returned to the home cage to rest for five minutes between each phase. Time spent with the novel and familiar objects during the test phase was measured using ANY-maze 7.00 software and the difference between the two for each subject was analyzed.

### Beam Walk

Between days 8-9, rats were trained for motor behavior tasks. A detailed description of the balance beam has been published^36, 38^. We used a tapered balance beam equipped with sensors detecting foot faults across all three segments (wide, middle, thin) to assess fine motor coordination. Animals were acclimated over two days with three training trials per day. During training, they were placed in a goal box for 60 seconds, then on a start platform to cross the beam. Testing occurred over three trials on day 10 with identical conditions. Foot faults and goal box latency were recorded and averaged for each subject.

### Rotarod

The Rotarod test is commonly used in Parkinson’s disease models to assesses equilibrium and motor function using a 4 cm diameter rotating rod that gradually increases in speed^38^. Animals were acclimated over days 8-9, with three training trials per day. During training, they were placed on the rod rotating at 5 rpm for 3 minutes, and if they fell, they were immediately returned to the rod. Testing occurred over three trials on day 10. The rod started at 1 rpm, accelerating at a rate of 0.2 rpm/s to a maximum of 50 rpm over 245 seconds. The time before animals fell off was recorded and averaged across trials.

### Magnetic Resonance Imaging

Imaging was conducted at two timepoints: day 3 (1-2 hours post-injury and treatment) and day 22 (three weeks post-injury and treatment). During the first session, rats were anesthetized with 0.5-1% isoflurane for structural T2-weighted and diffusion weighted imaging (DWI). The entire anesthetized imaging protocol lasted approximately 60 minutes (10 min setup, 6 min T2-weighted MRI, 44 min DWI). For a week before the second session, rats were acclimated for awake imaging. The second session included awake T2-weighted imaging, resting-state functional connectivity (rsFC), and functional MRI with hypercapnic challenge (fMRI), and concluded with anesthetized DWI. The entire imaging protocol lasted approximately 90 minutes, including 45 minutes awake (10 min setup, 6 min T2-weighted MRI, 15 min rsFC, 15 min fMRI) and 45 minutes anesthetized with 0.5-1% isoflurane (DWI).

Imaging sessions were conducted under dim red-light illumination using a Bruker Biospec 7.0 T/20-cm USR horizontal magnet (Bruker, Billerica, MA, USA). Both imaging sessions began with acquisition of a high-resolution T2-weighted anatomical data set using a rapid acquisition, relaxation enhancement (RARE) pulse sequence to screen for motion and to ensure there was no skull fracture or neuroanatomical injury as a result of rmTBI. Due to the length of the imaging procedure, the study was conducted in six staggered cohorts of four rats, with each experimental group randomly distributed across all six cohorts. The animal setup and imaging protocol has been described in detail in previous publications^33, 39–41^.

### Diffusion Weighted Imaging

DWI was acquired with a 3D spin-echo echo-planar-imaging (3D-EPI) pulse sequence having the following parameters: TR/TE = 500/20 ms, eight EPI segments, and 10 non-collinear gradient directions with a single b-value shell at 1000s/mm^2^ and one image with a B-value of 0 s/mm^2^ (referred to as B0) as previously described^24, 26, 42^. Geometrical parameters were: 48 coronal slices, each 0.313 mm thick (brain volume) and with in-plane resolution of 0.313 x 0.313 mm^2^ (matrix size 96 x 96; FOV 30 mm^3^). The imaging protocol was repeated two times for signal averaging. For statistical comparisons among rats, each brain volume was registered to the 3D MRI rat brain atlas for generation of voxel- and region-based statistics. All image transformations and statistical analyses were carried out using the in-house EVA software (Ekam Solutions LLC, Boston, MA, USA). The average value for each region of interest was computed using map files for indices of apparent diffusion coefficient (ADC) and fractional anisotropy (FA). Statistical differences in measures of DWI between experimental groups were determined using a nonparametric Kruskal Wallis multiple comparisons test (critical value set at <0.05) followed by post hoc analyses using a Wilcoxon rank-sum test for individual differences.

### Acclimation for awake imaging

To mitigate the stress associated with head restraint, rats were acclimated to the restraining and imaging protocol as previously described^24, 43^. Acclimation sessions were conducted daily for five consecutive days, progressively increasing in duration up to 45 minutes, the length of the awake imaging setup and acquisition protocol. Respiration, heart rate, motor activity, and plasma corticosterone levels significantly decrease over the course of the acclimation period^44^. This reduction in autonomic and somatic signs of arousal and stress improves signal resolution and image quality.

### Resting-State Functional Connectivity

A detailed description of the data acquisition, preprocessing, registration, and analysis has been previously described^33, 41, 45^. Scans were collected using a spin-echo triple-shot EPI sequence with the following parameters: matrix size = 96 x 96 x 20 (H x W x D), TR/TE = 1000/15 ms, voxel size = 0.312 x 0.312 x 1.2mm, slice thickness = 1.2 mm, 200 repetitions, time of acquisition 15 min. Image preprocessing combined AFNI, FSL, DRAMMS, and MATLAB software. After manual skull stripping, data underwent motion correction, outlier removal, slice timing correction, and affine registration to the rat atlas using DRAMMS. Data were then band-pass filtered (0.01-0.1 Hz), detrended, and smoothed (FWHM = 0.8 mm). Nuisance regression removed motion parameters, white matter, and CSF signals. Network analysis was performed in Gephi software using undirected networks from absolute connectivity matrices. Degree centrality was calculated as the sum of connections between each node and all other nodes.

Statistical analysis used GraphPad Prism 10.0, with Shapiro-Wilk tests determining normality. Paired t-tests or Wilcoxon signed-rank tests (for non-normal data) compared degree centrality between groups.

### Functional MRI with Hypercapnic Challenge

Functional images were captured using a multi-slice Half-Fourier Acquisition Single-Shot Turbo Spin Echo (HASTE) pulse sequence with an in-plane resolution of 312.5 μm². The scanning session for CO2 challenge lasted 15 minutes with 10 acquisitions per minute. Each scanning session was continuous, starting with a 5-minute baseline followed by 5 minutes of 5% CO2 exposure and 5 minutes after cessation of CO2.

A detailed description of the data analysis for functional changes in BOLD signal following CO2 challenge is published^23, 46^. In brief, the fMRI data analysis consisted of three main steps: pre-processing, processing, and post-processing. All these steps were executed using SPM-12 (available at https://www.fil.ion.ucl.ac.uk/spm/) and in-house MATLAB software. In the pre-processing stage, several operations were performed, including co-registration, motion correction, smoothing, and detrending. In the processing stage, all images were aligned and registered to a 3D Rat Brain Atlas©, which included 173 segmented and annotated brain regions. This alignment was performed using the GUI-based MIVA software developed by Ekam Solutions LLC (Boston, MA, USA). All spatial transformations applied were compiled into a matrix for each subject. Each transformed anatomical pixel location was tagged with its corresponding brain area, resulting in fully segmented representations of individual subjects for localization of functional imaging data to precise 3D volumes of interest.

Each scanning session consisted of 150 data acquisitions. The average signal intensity in each voxel of the first 5 min of baseline (acquisitions 1–50) was compared to 5–10 min (acquisitions 51–100) of CO2 exposure. We refer to the number of voxels in each brain area that showed a significant increase in BOLD signal above threshold as volume of activation. The mean volume of activation is the average of all rats for each experimental condition for that brain area. Statistical t-tests were performed on each voxel (∼ 36,000 voxels in the whole brain) of each subject within their original coordinate system. The baseline threshold was set at 1%. The t-test statistics used a 95% confidence level (p<0.05), two-tailed distributions, and heteroscedastic variance assumptions. As a result of the multiple t-test analyses performed, a false-positive detection controlling mechanism was introduced using the formula P(i)≤(i/V)(q/c(V)) across 173 brain areas, where q=0.2 and c(V)=1.

### Lipid Extraction and Partial Purification of plasma

Blood samples were collected 30 minutes after dosing on day 3 via the lateral tail vein. Plasma was isolated via centrifugation and stored in 75µL samples at -80°C until processed as previously described^47, 48^. In brief, methanolic extracts were partially purified using C^18^ solid phase extraction columns (Agilent, Santa Clara, CA, USA). Final elutions (i.e. fractions) of 65, 75, and 100 percent methanol were collected and stored at -80°C until mass spectrometry (MS) analysis.

### Lipidomics analysis

Methanolic elutions were analyzed as previously described^49^ with the exception that the API 7500 (Sciex, Framingham, MA 01701, USA) was used for analysis instead of the API 3000. The API 7500 is coupled to a Shimadzu LC system LC-40DX3 (Kyoto, Japan). Standard curves were generated by using purchased standards (Cayman Chemical, Ann Arbor, MI, USA), and those made in-house were validated through NMR and MS analysis as previously described^50^. Sciex Analyst peak matching software (Sciex, Framingham, MA 01701, USA) was used to validate standard peaks and sample peaks.

Statistical analyses for the plasma lipids were completed in IBM SPSS Statistics 29 (Chicago, IL, USA). One-way ANOVAs followed by Fisher’s Least Significant Analysis of individual endogenous lipids in plasma for each experimental group were analyzed using Student’s t-tests set to 2-tails and Type 2. Samples with an endogenous lipid concentration outside of 2 standard deviations from the group mean were omitted from statistics for that compound. Statistical significance for all tests was set at p < 0.05, and trending significance at 0.05 < p < 0.10. Descriptive and inferential statistics were used to create heatmaps for visualizing changes in the concentration of each lipid analyte for every condition as previously described^51^.

### Molecular biology, Western blot, and solubility fractionation

On day 23, rats were deeply anesthetized via isoflurane inhalation for tissue collection. Brains were rapidly extracted after decapitation, frozen, and stored at -80°C until proteomic analysis. Rat brain samples of frontal area were isolated and solubilized with Douce homogenizer followed by sonification in standard RIPA (50 mM Tris pH 8, 150 mM NaCl, 0.5% sodium deoxycholate, 0.1% SDS and 1% NP40) with protease/phosphatase inhibitor as previously described^52^. The solubility fractionation was modified and 200μg total protein from the RIPA-soluble fraction was centrifuged at 180,000g for 30 minutes, the supernatant was removed, and the RIPA-insoluble pellet was washed in RIPA buffer and solubilized in 7M urea/2M thiourea before Western blot as previously described^53^. RIPA samples of 20μg each were run on a 4-20% SDS-PAGE and transferred to PVDF, blocked, probed, and visualized as previously described (antibody table)^52^.

### Statistical analysis for molecular biology

An ANOVA with Tukey post hoc was performed comparing each of the three groups (*n* = 8 biological replicates/group) reported as mean ± SD. All blots were exposed under the same conditions. All raw uncropped data are available upon request.

## Results

### Behavior

Shown at the top of **Fig 1a** is a timeline for experimental procedures and data collection. Shown in **Fig 1bc** are the results from the four behavioral assays: Open Field, Beam Walk, Rotarod and Novel Object Recognition. The data collected from each assay is organized into Motor Behavior (top row) and Cognitive and Emotional Behaviors (lower row). The data are shown as dot plots (subjects) and bar graphs (mean ± 95% confidence interval) and analyzed using a one-way ANOVA. HTR was collected each of day 1-3 immediately following PSI or VEH treatment with the other four assays conducted between day 4 and day 10. HTR was observed in all PSI-treated rats exclusively, but only after the first dose and not after subsequent administration on day 2 and day 3. This adaptation has been observed in rodents chronically exposed to hallucinogenic 5-HT2A agonists^54–56^. No significant motor differences were observed across groups for the Beam Walk (time to reach the goal box and number of slips), Rotarod (latency to fall), or Open Field (total distance traveled and average speed).

Episodic learning and short-term memory were not significantly different between groups when tested for object recognition. Measures of anxiety associated with fear of open spaces (Agoraphobia) or attachment to protected surfaces (Thigmotaxis) were also not significantly different between experimental groups in the Open Field.

### Head Injury and Diffusion Weighted Imaging

Shown in **Fig 2a** are radiograms of each subject for the three experimental groups characterizing the head injury associated with the site of impact near the prefrontal cortex. The areas of impact are characterized by swelling and edema to the soft tissue over the skull, visualized as white contrast in T2-weighted imaging (indicated by arrows). Importantly, no skull fracture or contusion is observed. In three-week follow-up scans, all impact-site edema is resolved. **Fig 2b** reports changes in ADC values collected during DWI 1-2 hours post-injury and treatment on day 3 as a proxy for brain edema. The box insert shows data from the whole brain (173 brain areas). There is a pronounced increase in whole-brain ADC values (paired t-test, p<0.0001) comparing SHAM-VEH and rmTBI-VEH rats with a mean difference of 0.06. When comparing all three experimental groups (matched one-way ANOVA), both SHAM-VEH controls and rmTBI-PSI treated rats were significantly less than rmTBI-VEH untreated rats (p<0.0001). However, it should be noted that PSI treatment did not reduce all of the edema, as whole-brain measures were still significantly greater than SHAM-VEH. When the 173 brain areas in the rat atlas are organized into brain regions, it is possible to delineate those areas that are more or less sensitive to head injury and PSI treatment as measured by changes in ADC. These data are shown in the bar graphs (mean ± SD) and dot plots (subregions). For example, the hippocampus, comprised of nine brain subregions, showed the same whole-brain profile of ADC values with SHAM-VEH controls and rmTBI-PSI treated rats being less than rmTBI-VEH untreated rats, but with rmTBI-PSI still greater than SHAM-VEH. This is also true for the somatosensory cortex (SS cortex) and the prefrontal cortex. PSI treatment reduced edema to SHAM-VEH levels in the basal ganglia, thalamus, cerebellum, and olfactory system. Specifically, an inverse ADC:FA correlation is detected, indicating whole-brain vasogenic edema. On three-week follow-up scans, all ADC and FA differences were resolved; however, lasting functional changes were detected.

**Fig 2.**
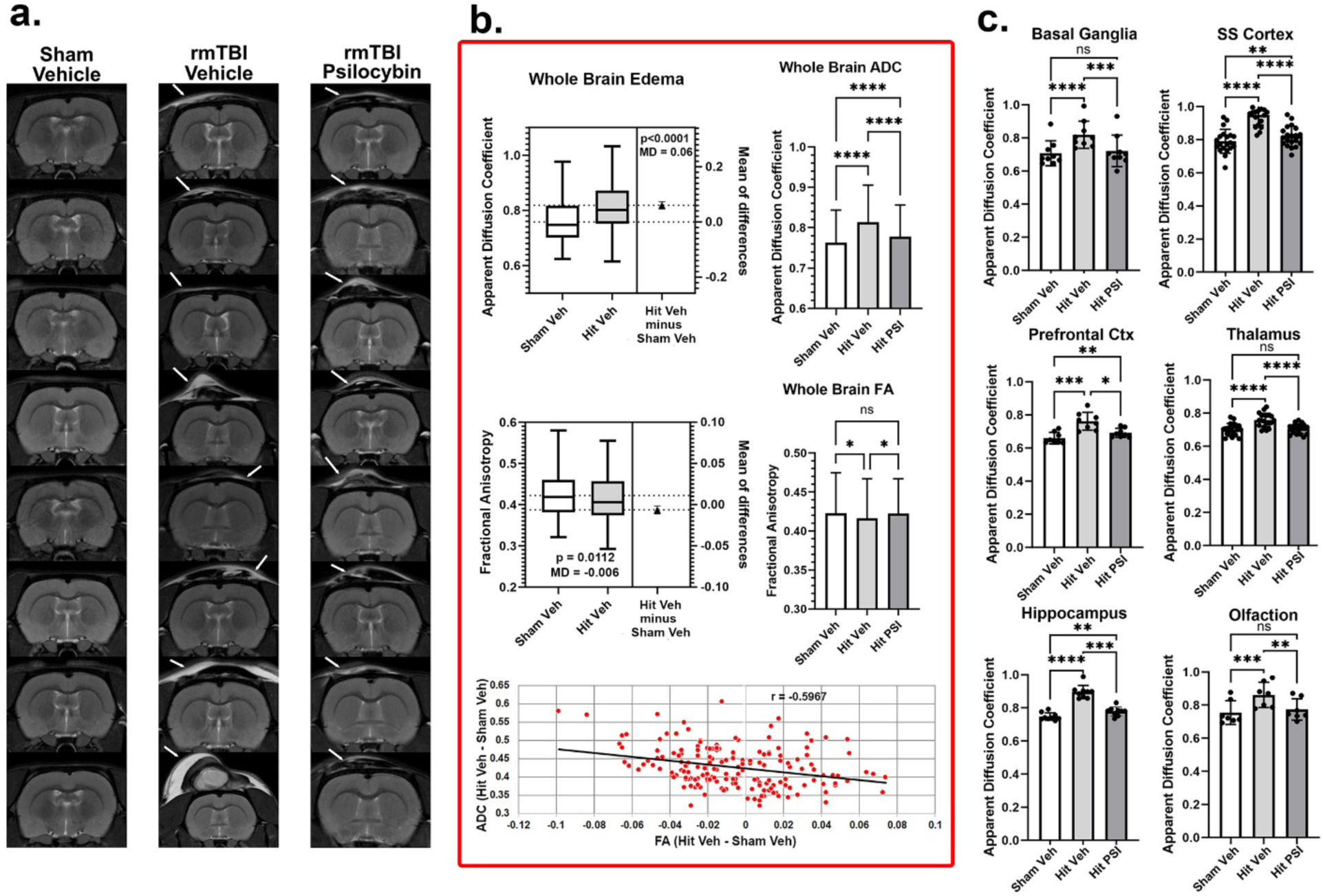
Diffusion Weighted Imaging: Vasogenic edema reduced by PSI. T2-Weighted Imaging: **a.**) Shown are radiograms of frontal sections of the brain of all subjects taken following the last of three head impacts. The arrows point to the approximate site of impact identified by T2-weighted enhanced signal showing edema on the skin above the skull, but no skull fracture or contusion. This is obvious in all hit rats but not in sham controls. Diffusion Weighted Imaging: **b.**) Shown are whole-brain changes in water diffusivity as measured by apparent diffusion coefficient (ADC) and fractional anisotropy (FA). When comparing SHAM-VEH and rmTBI-VEH there was a global increase in ADC and decrease in FA. There was a significant inverse relationship between ADC and FA values (r = -.5967) as shown in the bottom graph. Each red dot is a brain area taken from the rat 3D brain atlas. A comparison across all three experimental groups shows that PSI treatment significantly reduces ADC and increases FA values, reversing the effects of head injury. **c.**) Shown are the regional changes in ADC values for each of the experimental groups. In all brain regions, rmTBI-VEH rats show a significant increase over SHAM-VEH controls, while rmTBI-PSI treated rats show a significant decrease compared to rmTBI-VEH rats. Error bars show standard deviation. * p<0.05, ** p<0.01, *** p<0.001, **** p<0.0001

### Vascular Reactivity to Hypercapnia

**Fig 3** shows the change in vascular reactivity to carbon dioxide (CO2) challenge three weeks post-injury and treatment. The box insert shows the number of voxels activated for positive BOLD for the whole brain for each of the experimental conditions. A matched one-way ANOVA shows a significant difference between groups (F(1.583, 267.5) = 90.44, p<0.0001). rmTBI-VEH untreated rats present with significantly higher (p<0.0001) voxel numbers than SHAM-VEH controls or rmTBI-PSI treated rats (Tukey’s multiple comparison post hoc test), indicating vascular hyperreactivity. The bar graphs (mean ± SD) and dot plots (subregions) show the regional differences in vascular reactivity. In all cases, a matched one-way ANOVA showed a significant difference between groups. Post hoc tests showed that rmTBI-VEH untreated rats were significantly higher than SHAM-VEH controls and, in the case of the basal ganglia, prefrontal cortex, and olfactory system, significantly greater than rmTBI-PSI treated rats. It should be noted rmTBI-PSI treated rats still showed values significantly higher (p<0.0001) than SHAM-VEH controls in all brain regions with the exception of the prefrontal cortex, suggesting the treatment was not totally effective in reducing the hyperreactivity to CO2 challenge following head injury.

**Fig 3.**
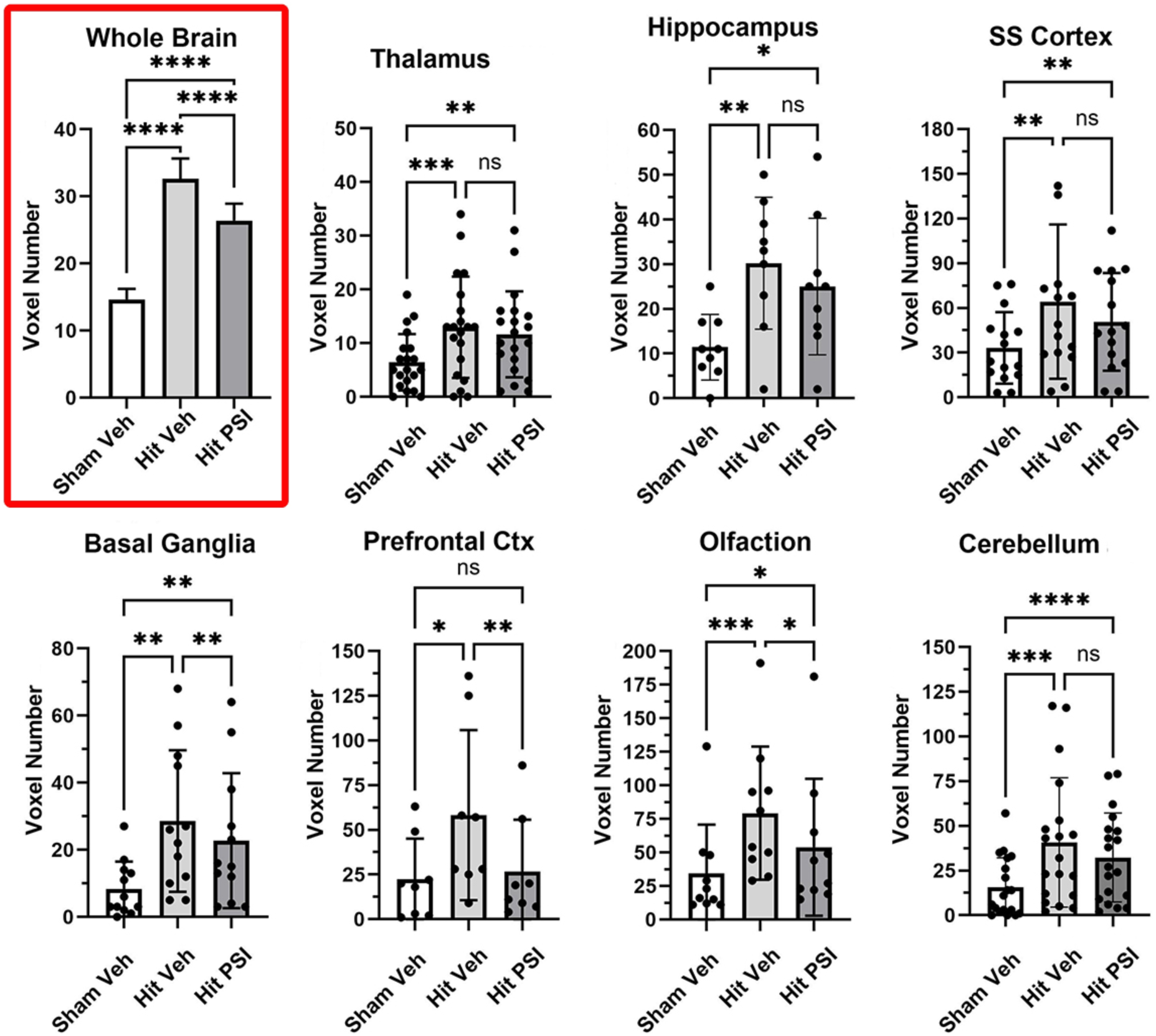
Hypercapnic Challenge: Vascular hyperreactivity reduced by PSI. Shown are changes in vascular reactivity (positive BOLD voxel number) in response to a 5% CO2 challenge. Whole-brain (box) was significantly elevated above control in rmTBI-VEH rats (gray bars). rmTBI-PSI treated rats were significantly lower than rmTBI-VEH untreated rats. Regional differences in vascular reactivity are shown in the bar and dot plots (mean ± SD). Each dot is a subregion in that brain region. * p<0.05, ** p<0.01, *** p<0.001, **** p<0.0001

### Resting-State Functional Connectivity

The bar graphs (mean ± SD) in the highlighted box of **Fig 4a** show the connections to the whole brain for all 173 brain areas three weeks post-injury and treatment. The global statistics using graph theory analysis for the SHAM-VEH controls were Average Degree 14.737, Graph Density 0.087 and Average Path Length 2.541. For rmTBI-VEH untreated rats, the statistics were Average Degree 9.088, Graph Density 0.053, and Average Path Length 2.909. For rmTBI-PSI treated rats, the Average Degree was 35.532, Graph Density 0.209, and Average Path Length 1.892. There was a significant treatment effect using a matched one-way ANOVA [F(1.352,229.8) = 493.6, p<0.0001)] followed by Tukey’s multiple comparison post hoc test (**** p<0.0001). The significant differences between each experimental group for different brain regions are shown as dot and bar graphs. The dots represent the different subregions within each brain region.

**Fig 4.**
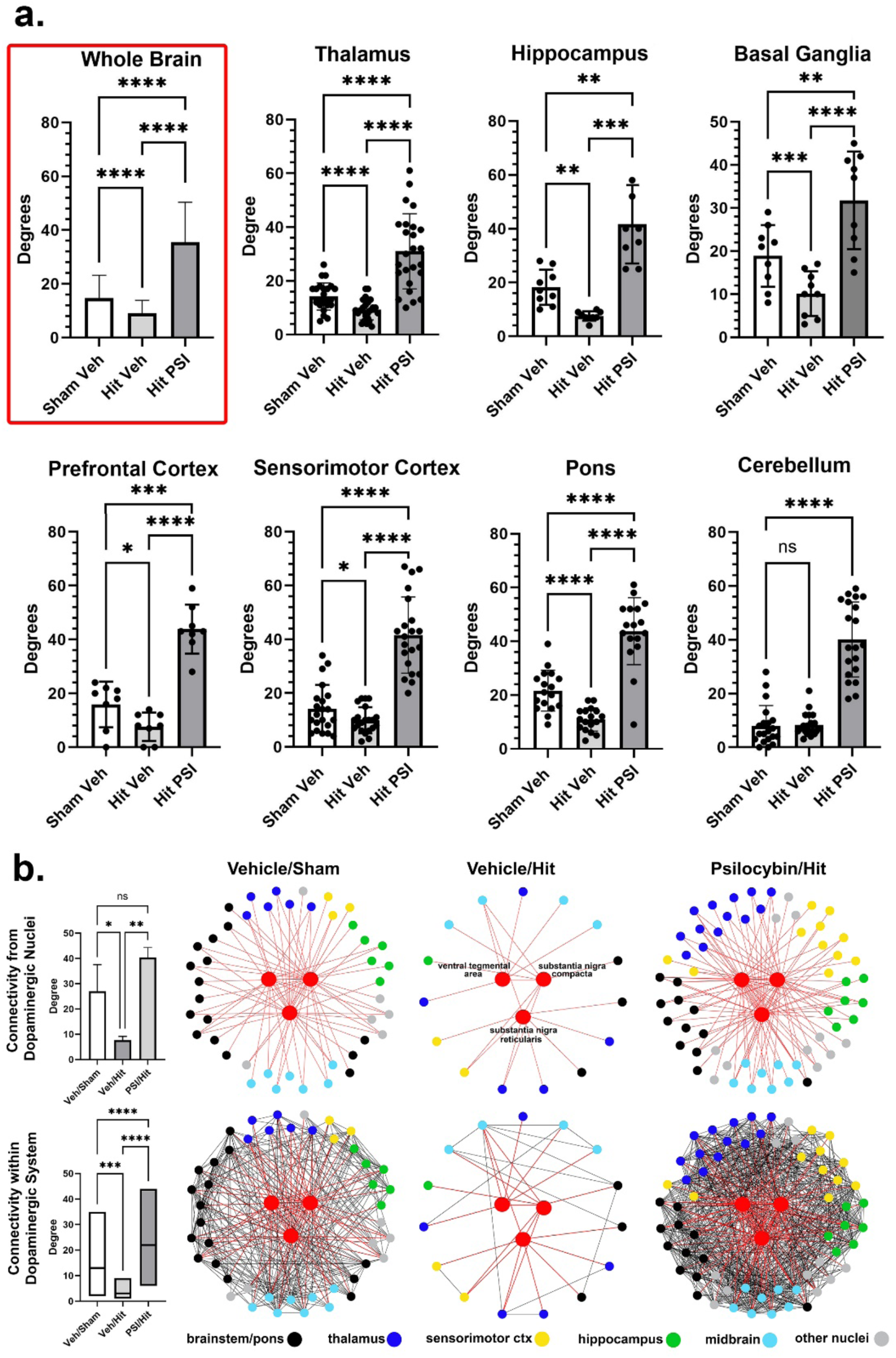
Functional Connectivity: Hypoconnectivity reversed by PSI. **a.**) Shown are the number of degrees (connections) for the whole brain (red box) for each experimental group (mean ± SD). Regional differences in degrees are shown in the bar and dot plots (mean ± SD). Each dot is a subregion of that brain region. In all cases, rmTBI-PSI treated rats showed greater connectivity than SHAM-VEH or rmTBI-VEH rats. **b.**) The connectivity to the dopaminergic system for each experimental condition is shown. The top bar graphs (mean ± SD) to the left show the number of efferent connections from the three major dopaminergic nuclei in the midbrain (ventral tegmental area, substantial nigra compacta, and substantia nigra reticularis) while the radial connectivity maps to the right depict these connections to color-coded brain regions. The bottom bar graphs to the left (median and 1^st^ and 3^rd^ quartile) show the number of connections between and within the dopaminergic system. * p<0.05, ** p<0.01, *** p<0.001, **** p<0.0001

**Fig 4b** shows changes in connectivity to the three midbrain dopaminergic nuclei (ventral tegmental area, substantia nigra compacta, and substantia nigra reticularis). The panels above show the nodes (colored circles) and connections, or edges, (red lines) from the three key dopaminergic nuclei (red circles). This functional connectivity is displayed for all three experimental groups. The differences are dramatic, as rmTBI-VEH untreated rats show an extreme loss of connectivity as compared to SHAM-VEH controls and rmTBI-PSI treated rats. Psilocybin promotes hyperconnectivity, recruiting nodes and connections to and within the thalamus (blue circles) and sensorimotor cortices (yellow circles). The network connections between all nodes (black lines) are shown in the lower panels. rmTBI-VEH untreated rats have very few connections, while the network connectivity in SHAM-VEH controls, and especially rmTBI-PSI treated rats, is very pronounced. These differences in degrees, or all connections associated with the three dopaminergic nuclei, are shown in the top bar graph (mean ± SD). Both SHAM-VEH controls (p<0.05) and rmTBI-PSI treated rats (p<0.01) were significantly greater than rmTBI-VEH untreated rats (matched one-way ANOVA, F(1.062, 2.124) = 32.59, p = 0.025). The difference in degrees associated with the entire dopaminergic system is shown in the bottom bar graph (median, min, and max). Again, SHAM-VEH controls (p<0.001) and rmTBI-PSI treated rats (p<0.0001) were significantly greater than rmTBI-VEH untreated rats (Kruskal-Wallis test for non-parametric data p<0.0001). rmTBI-PSI treated rats were also greater than SHAM-VEH controls (p<0.0001), evidence of hyperconnectivity.

### Phosphorylated tau reduces to control levels after PSI treatment

The RIPA-soluble protein analysis normalized to tubulin shows a significant increase in phosphorylated tau (PHF-1, provided by Peter Davies) when comparing SHAM-VEH controls to rmTBI-VEH untreated rats (*p* = 0.0017). Interestingly, rmTBI-PSI treated rats show a reduction in phosphorylated tau back to near SHAM-VEH levels (*p* = 0.0015; **Fig 5ab**). The aggregated RIPA-insoluble (urea soluble) phosphorylated tau (PHF-1) shows a significant increase in rmTBI-VEH (*p* = 0.0318) but no change in rmTBI-PSI as compared to SHAM-VEH (*p* = 0.2654; **Fig 5cd**). Though there is no significant decrease in phosphorylated tau aggregation comparing rmTBI-VEH to rmTBI-PSI, there is a distinct clustered trend in reduced aggregated tau and no significant change when SHAM-VEH is compared to rmTBI-PSI (*p* = 0.5063). This lack of urea-soluble aggregation of tau may be due the time of evaluation after injury and future research to examine age-related changes after treatment may show that sustained aggregation may not develop with PSI treatment due to the initial reduction in RIPA-soluble tau. This reduction in aggregation is consistent to what was observed in the RIPA-soluble fraction. However, this significant increase in phosphorylated tau post-injury may be at an early pre-tangle aggregation stage that is reduced to normal levels in PSI-treated animals.

**Fig 5.**
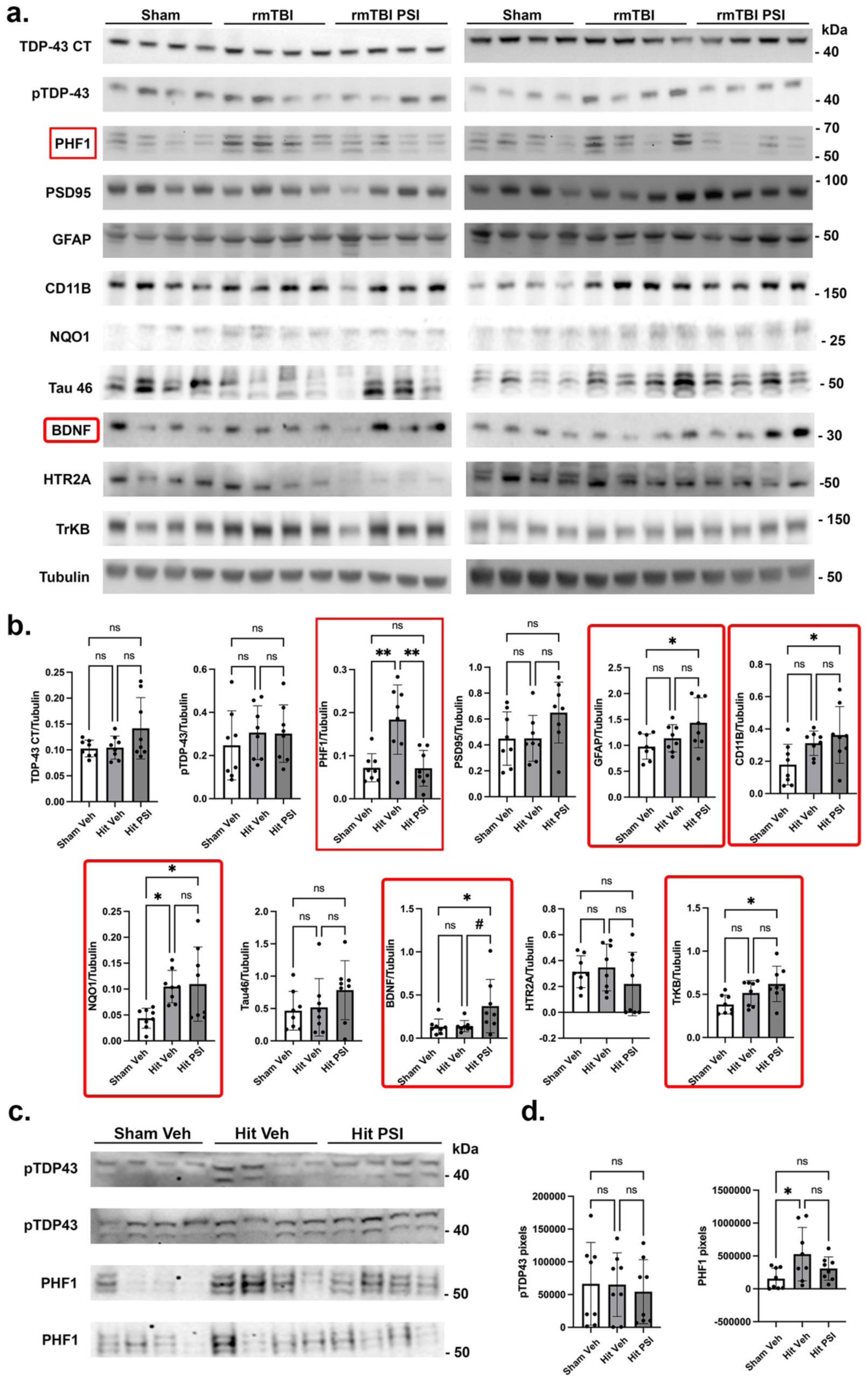
Proteomics: Phosphorylated tau reduced to control levels by PSI. **a.**) Western blot of RIPA-soluble proteins. **b.**) Quantitative analysis normalized to tubulin of RIPA fractions. **c.**) Western blot RIPA insoluble/urea soluble fractions. **d.**) Quantitative analysis normalized to total protein. Statistical analysis: one way ANOVA with Tukey post hoc (*n* = 8 biological replicates/3 groups) (e.) * p<0.05 and ** p<0.002 (all p vales see table).

In addition, GFAP (astroglia; *p* = 0.0378) and CD11b (microglia; *p* = 0.0294) levels are significantly increased in rmTBI-PSI treated rats relative to SHAM-VEH controls. Changes in gliosis may initially be a protective response to the injury, though it is not significantly different between SHAM-VEH controls and rmTBI-VEH untreated rats^57^ (**Fig 5ab**). When astrocyte activation is reduced in an APP/PS1 mouse model, accelerated age-related plaque deposition occurs which implies the need for functional astrocytes to respond to stress^58^. NAD(P)H quinone dehydrogenase 1 (NQO1), induced by the NRF2 antioxidant response pathway (review^59^), has been shown to protect against oxidative stress. rmTBI-PSI treated animals show no significant differences from rmTBI-VEH untreated rats, though there is an associated response increasing NQO1 in rmTBI-VEH (*p* = 0.0405) and rmTBI-PSI (*p* = 0.0260) groups relative to SHAM-VEH (**Fig 5ab**). In rmTBI-PSI treated rats, BDNF shows a significant increase over SHAM-VEH controls (*p* = 0.0486) and a clustered increase over rmTBI-VEH untreated rats (*p* = 0.0569). BDNF exerts its effects via the high-affinity tyrosine kinase receptor TrkB^60^.

Interestingly, TrkB (NTRK2) shows a significant increase in rmTBI-PSI treated rats (*p* = 0.0168) compared to SHAM-VEH controls, which may be a protective measure increasing downstream factors (**Fig 5ab**). No significant changes were observed among other RIPA-soluble proteins or urea-soluble phosphorylated TDP-43. These samples were from the frontal area, and the hippocampus and cerebellum may show additional modulations of these pathways. This variability in aggregation may be consistent with the experimental dynamics of our rat rmTBI model consistent with the variability of the human disease. Though phosphorylated TDP-43 in some rats show distinct changes in aggregation, further evaluation of TDP-43, SNCA, amyloid, and other aggregated proteins will be investigated.

### Plasma signaling lipids

Two levels of analysis reveal important systemic effects of the intersection of our rmTBI model and repeated PSI treatment. **Fig 6** shows the comparisons of SHAM-VEH versus rmTBI-VEH where the heatmap illustrates the direction of change with rmTBI. Here we see that, overall, there were only three changes in circulating levels of signaling lipids; however, all changes were significant decreases after rmTBI (e.g. stearoyl leucine, stearoyl taurine, and palmitoyl taurine). By contrast, animals treated with PSI who underwent rmTBI had six times as many lipids change as a result of the treatment (3 versus 18 respectively), and all significant changes were increases. Key findings were that PSI treatment caused significant increases in most free fatty acids measured and caused reversal of levels of stearoyl taurine, suggesting that this signaling lipid may play a key role in effects of rmTBI.

**Fig 6.**
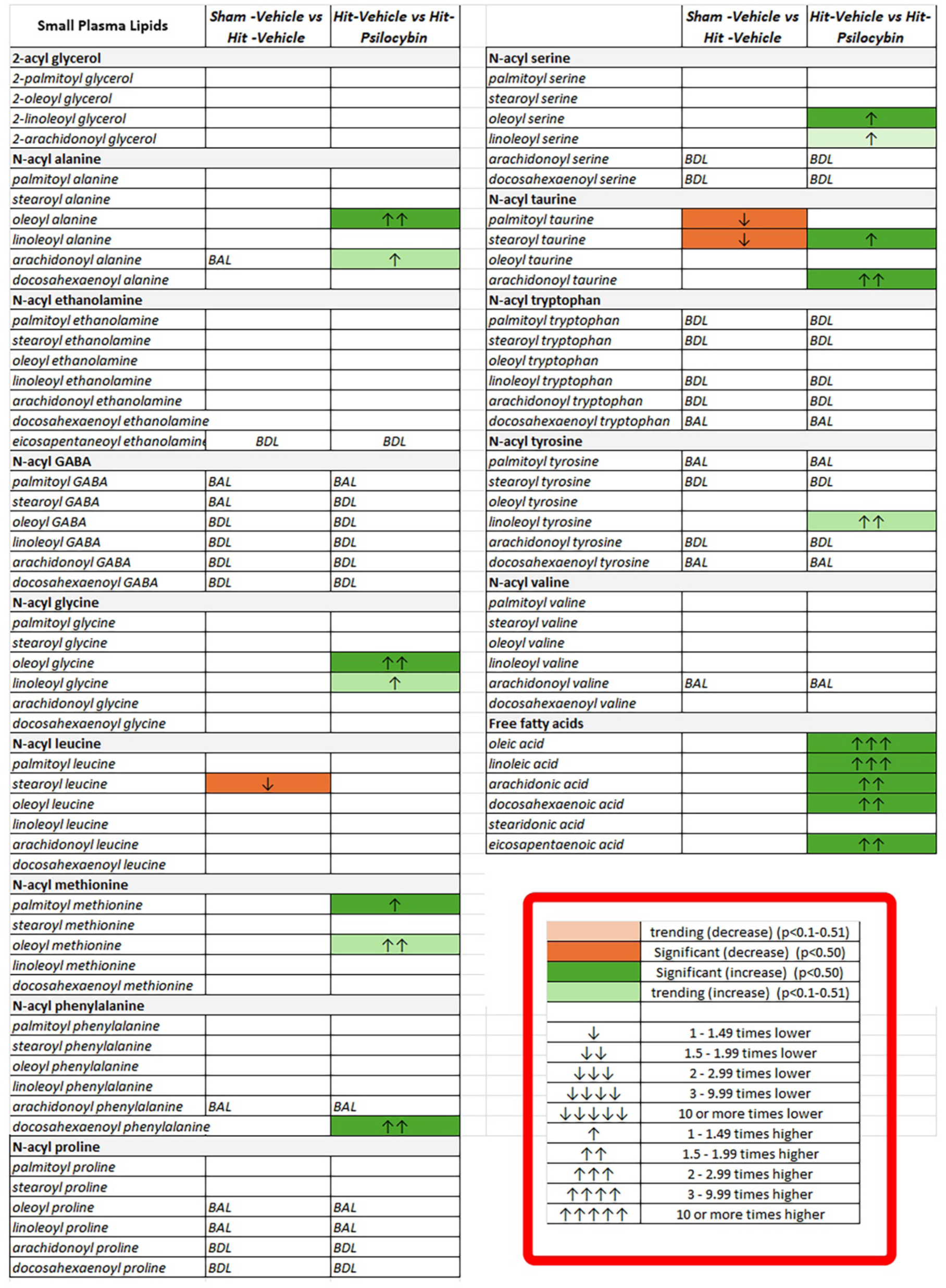
Lipidomics: Novel plasma lipid biomarkers modulated by rmTBI and PSI. The direction of changes for each analysis group relative to SHAM-VEH are depicted by color and arrow direction, with an increase represented by a green box and upward arrows and a decrease represented by an orange box with downward arrows (boxed insert). Level of significance is shown by color shade, wherein p < 0.05 is a dark shade and 0.05 < p < 0.1 is a light shade. Effect size is represented by the number of arrows, where 1 arrow corresponds to 1–1.49-fold difference, 2 arrows to a 1.5–1.99-fold difference, 3 arrows to a 2–2.99-fold difference, 4 arrows a 3–9.99-fold difference, and 5 arrows a difference of tenfold or more^126^. An abbreviation of ‘BDL’ indicates that the lipid concentration that was present in the sample was below the detectable levels of our equipment while ‘BAL’ indicates below analytical levels.

## Discussion

Repetitive mild TBI is very common and can occur over the life span, affecting adolescents playing organized sports, professional athletes, individuals in service, and the elderly. rmTBI is a significant risk factor for dementia, Parkinson’s, and Alzheimer’s^61^. Neuroinflammation, alterations in gray matter microarchitecture, impaired cerebral blood flow, disruption in blood-brain barrier (BBB) permeability, and dysfunction in clearance of unwanted phosphorylated proteins lie at the heart of these neurodegenerative diseases^62, 63^. The repetitive mild head injury used in this study was designed to reflect the human experience. Our neuroradiological findings recapitulate much of the neuropathology observed and measured in the clinic using MRI. Below, we discuss the “bench to bedside” relevance of these data and the significance of psilocybin as a putative treatment for head injury and neurodegenerative disease.

### Model of Repetitive Mild Head Injury

The guidelines from the Centers for Disease Control and Prevention, World Health Organization, and American Congress of Rehabilitation Medicine for diagnosing mild head injuries include self-reports of transient confusion, disorientation, impaired consciousness, or dysfunction in memory around the time of the injury with no neuroradiological evidence of structural damage to the brain^64, 65^. To that end, our lab adopted a head injury model in rats originally developed by Viano and colleagues^29^ and further refined by Mychasiuk et al.^30^, controlling for the axis of injury, rotational force, and head acceleration in different directions. We have added to this model by performing the mild head impacts while rats are fully awake without the confound of anesthesia and during the active period of their circadian cycle. As was the case in previous studies using this model^23, 66, 67^, all rats showed normal ambulatory behavior within seconds of being placed into their home cage after head impact. With this model there are no mortalities and no evidence of skull damage or contusion as determined by MRI. These are the “bump on head, ice pack” injuries that show up on radiograms as edema in the skin at the site of impact as shown in **Fig 2a**. In this model, the neurobiological effects of a single head impact resolve within 24 hours^25^. However, two or more head impacts separated by 24 hours each have long-term consequences, as noted in the clinical literature^68, 69^. In recent publications using this model of rmTBI, our lab has reported a constellation of neurobiological and neurochemical changes in the brains of male and female rats. The pathology includes vasogenic edema^25^, altered vascular reactivity^23, 70^ and gray matter microstructure^26^, disruption in blood-brain barrier permeability^27^, reduction in perivascular clearance^24^, increases in microgliosis^24, 26^, altered astrocytic AQP4 expression and polarization^24^, changes in brain functional connectivity^26^, and decreased brain-derived neurotrophic factor (BDNF) expression^66^. In the present study we also report elevated levels of phosphorylated tau in the prefrontal cortex of rats weeks after rmTBI.

### Mild Repetitive Head Injury and Behavior

Measures of motor activity and coordination and learning and memory were not significantly different across experimental groups. This is not unexpected with mild head impacts in rodents. Anesthetized rats exposed to a single mild head impact^30, 71^ or anesthetized mice subjected to five mild head impacts spaced 24 hours apart^72^ exhibit minor balance and motor coordination deficits that resolve within a few days. Ren et al. reported that anesthetized mice experiencing a single mild impact show no cognitive changes but do display a decrease in motor performance on the rotarod, lasting up to 24 days^73^. In contrast, mice subjected to two mild, repetitive head impacts while fully awake, evaluated through a series of neurobehavioral tests, show complete recovery within hours^74^. Whether the early impacts in our study have long-term effects on behavior as the subjects age remains uncertain. Recurrent mild head injury in humans can lead to persistent post-concussive symptoms that overlap with other psychiatric disorders like PTSD and major depression^75^.

### Vasogenic Edema

Edema plays a major role in the neuropathology of head injuries^76, 77^. Vasogenic edema results from damage to the BBB, leading to the immediate movement of fluid into the brain’s extracellular space. An increase in apparent diffusion coefficient (ADC), which measures water mobility, serves as an indicator of this volume change^77^. This increase in ADC is typically associated with a decrease in fractional anisotropy (FA). In cases of moderate-to-severe head injury, cytotoxic edema occurs, marked by cellular swelling due to disrupted osmolarity regulation across the plasma membrane. This type of edema generally shows a decrease in ADC and an increase in FA^78^.

In a previous study we found that a single mild impact, which showed no neuroradiological evidence of brain damage, led to a temporary increase in vasogenic edema in the thalamus, basal ganglia, and cerebellum, as indicated by a rise in ADC^25^. This increase in extracellular fluid volume peaked at 6 hours and returned to baseline by 24 hours. In the present study using three mild head impacts, there was a global increase in ADC values suggestive of vasogenic edema. This increase was shared by several brain regions including the basal ganglia, hippocampus, thalamus, prefrontal cortex, and somatosensory cortex. In each of these cases, treatment with PSI significantly reduced the ADC values. In the basal ganglia and thalamus, these values were reduced to levels measured in SHAM-VEH controls. How does PSI reduce ADC? We would suggest two possible mechanisms: 1) Support BBB structural integrity via endothelial tight junctions to reduce vasogenic edema, and/or 2) enhance astrocytic activity helping to promote convection of excess extracellular fluid through aquaporin channels lining the astrocytic endfeet. With respect to the first, there have been no studies focused solely on PSI and capillary BBB integrity. However, there have been several studies following changes in gene transcription in the cortex in rodents in response to exposure to 5-HT2A psychedelics^79–82^. Most relevant to this study, Jefsen and coworkers showed a significant increase in transcription of select genes in the prefrontal cortex of rats in response to a single dose of PSI ^80^. Several of these genes, including Cebpb (CCAAT enhancer binding protein beta); Iκβ-α (NFKBIA), a key regulator of NF-κB signaling; and Nr4a1 (Nur77) are all involved in vascular inflammation and endothelial cell function.

In terms of enhanced fluid convention, we have hypothesized in a previous study^83^ that astrocytes localized around capillaries form a hydrolytic syncytium connected by gap junctions^84^. Free water from vasogenic edema would move down osmotic gradients, promoting swelling of the astrocytes and hydrostatic pressures favoring convection toward AQP4 water channels at the endfeet surrounding capillaries. PSI causes an elevation in GFAP (Glial Fibrillary Acidic Protein), a filament protein that forms a dynamic intracellular scaffold that interacts with other binding proteins like plectin (cytoskeletal crosslinker) and α-actinin (actin bundling protein) to form a mechanically integrated system that affects astrocytic cell volume^85^. Changes in astrocyte calcium fluctuations enhance phosphorylation and reorganization of the filament structure that could cause mechanical forces, combined with passive osmotic forces, to promote convection, lowering extracellular fluid and decreasing ADC values.

### Vascular Reactivity

Fundamental to brain health is autoregulation of cerebral blood flow in the face of fluctuations in systemic blood pressure. At the level of the neurovascular unit, homeostasis is maintained by local changes in vascular reactivity and capillary blood flow in response to the surrounding metabolic environment. A simple biomarker for evaluating the health of cerebral blood vessels is a hypercapnic challenge^86^ causing a passive expansion of blood vessels, decreased resistance, and heightened blood flow. Importantly, when exposed to higher levels of CO2, there is no change in metabolic oxygen consumption. Consequently, the alteration in the MRI signal caused by the increased blood flow in the brain is directly linked to the change in the partial pressure of CO2. The change in BOLD signal in response to heightened CO2 levels is a straightforward and reliable technique for evaluating the cerebral vascular reactivity (CVR) in functional imaging studies^87–89^. CO2-induced changes in BOLD have been used in the clinic to evaluate the health of cerebral vasculature in several neurological disorders including Alzheimer’s^90–92^, stroke^93^, multiple sclerosis^94^ and TBI^95^. A recent study by Liu and colleagues evaluated the reliability of hypercapnia-driven CVR as a biomarker for cerebrovascular function and found it was suitable across different scanning platforms and imaging sites for use in longitudinal studies and clinical trials^96^.

There have been numerous studies using BOLD imaging and hypercapnic challenge to evaluate cerebrovascular function following TBI. With moderate-to-severe TBI there is a significant decrease in CVR in response to CO2 challenge^95^. Subjects with serious cerebral vascular injury following TBI show reduced global CVR compared to healthy controls^97–99^. Reports of reduced CVR to CO2 challenge are also true in preclinical studies of TBI^100, 101^. However, there are reports of increased CO2-induced CVR following mild head injury, ^102^ particularly in those with sports-related concussions accompanied by reduced CBF and cerebral metabolism^103, 104^. This would suggest enhanced regulation of blood flow to affected areas with reduced metabolism^103, 104^. In this and a previous study on repetitive mild head injury in female rats, ^23^ we found an increase in CVR with CO2 challenge. PSI treatment significantly reduced the increase in whole-brain CVR as compared to rmTBI-VEH untreated rats but was still significantly elevated above SHAM-VEH controls. The basal ganglia, prefrontal cortex, and olfactory system were significantly reduced with PSI treatment as compared to rmTBI-VEH untreated rats.

### Functional Connectivity

This model of rmTBI presents with global functional hypoconnectivity. This result is not unexpected given the many reports in clinical studies showing a similar decrease in connectivity following head injury. Patients studied within the first seven days of a mild TBI and presenting with post concussive syndrome (PCS) show a reduction in functional connectivity in the sensorimotor and central executive networks as compared to healthy volunteers^105^. Patients with mild TBI and PCS also show decreases in connectivity to the thalamus^106^ and a decrease in the symmetry of connectivity between left and right thalamic nuclei^107^. The connectivity between the motor-striatal-thalamic network is also reduced in mild TBI while the frontoparietal network is increased^108^.

Pinky et al. reported a decrease in functional connectivity in the cerebellum and basal ganglia in sports-related concussions in 12-18-year-olds^109^. Most recently, Fitzgerald and colleagues ran longitudinal studies on young athletes competing in American football, collecting functional connectivity prior to, during, and following the season^110^. Patterns of functional connectivity declined during the playing season but recovered after the season ended. They proposed that exposure to multiple head acceleration events may cause changes in brain neurobiology that are similar to concussion but in the absence of any symptomatology. The most robust finding in the present study was the dramatic hyperconnectivity that followed treatment with PSI. Not only did PSI prevent the hypoconnectivity of head injury, it exceeded the normal connectivity of sham controls. This global phenomenon was consistent across multiple brain regions.

Given the fact that head injury is one of the major risk factors for the development of Parkinson’s disease, we focused on the connectivity of the midbrain dopaminergic system: the ventral tegmental area (VTA) and the substantia nigra (SN) compacta and reticularis. The efferent connections from these brain areas were significantly increased with PSI over rmTBI-VEH untreated rats with a shift toward greater connectivity to the thalamus and somatosensory cortex when compared to SHAM-VEH controls. The dopaminergic system, as defined by the connectivity within and between all these efferent targets from the VTA and SN, was significantly increased over SHAM-VEH and rmTBI-VEH untreated rats.

### Psilocybin, BDNF, TrkB, and Phosphorylated Tau

In a recent study using our head injury model, we reported decreased BDNF levels in the hippocampus and around the midbrain dopaminergic nuclei^66^. Both male and female rats 9 months of age show a significant decrease in BDNF in the hippocampus with head injury while males also showed a decrease in BDNF in the substantia nigra. BDNF is the most prevalent neurotrophin in the brain, crucial for the survival, differentiation, synaptic plasticity, and axonal growth of both peripheral and central neurons during adulthood^111^. BDNF exerts its effects via the high-affinity tyrosine kinase receptor TrkB ^60^. After spinal cord injury or TBI there is an increase in TrkB mRNA expression at the site of injury^112^. BDNF has been recognized for its significant involvement in cellular processes related to recovery after TBI, such as promoting neuronal survival, axonal growth, and the formation of new synapses^113^. Given its crucial role, BDNF has been extensively studied in experimental TBI models^114^. In a recent study, Moliner et al. reported PSI can directly bind to TrkB receptor with a very high affinity^115^. Further, Zhao et al recently demonstrated the ability of PSI to restore decreases in prefrontal cortex BDNF expression in a rodent model of major depressive disorder^116^. Here, we show that PSI elevated BDNF and TrkB proteins above SHAM-VEH control and rmTBI-VEH untreated rats in the prefrontal cortex. These data provide a possible mechanism of action for the healing effects of PSI.

Interestingly, the BDNF/TRK2 pathway is altered in neurodegeneration associated with tauopathies^117^, a group of neurodegenerative diseases characterized by the hyperphosphorylation of tau^118^. Tau proteins bind to and stabilize microtubules in neurons to help maintain the structure and function of axons^119^. When tau becomes hyperphosphorylated, it can aggregate into neurofibrillary tangles (NFTs) inside neurons, disrupting cell function^120^. The accumulation of NFTs and the subsequent loss of neuronal function are key features in Alzheimer’s disease, frontotemporal dementia, and progressive supranuclear palsy^121–123^. Here, we provide evidence that PSI can reduce phosphorylated tau following head injury, which may ultimately reduce the risk for neurodegenerative diseases.

### Plasma lipids as biomarkers of CNS effects

Recently we showed that injections of the endogenous signaling lipid, palmitoyl ethanolamine (PEA) causes significant changes in CNS connectivity and behavior as well as plasma and CNS signaling lipids^33^. These data demonstrate that the presence of signaling lipids in plasma have a direct effect on CNS activity. Data here show that rmTBI alone has a modest but significant effect on signaling lipids by significantly reducing some of the lipids. By contrast, PSI treatment, which significantly improved rmTBI outcomes at multiple levels, drove significant increases in a wide range of circulating signaling lipids. In both cases, these signaling lipids may be key biomarkers for both injury and recovery.

The dramatic hyperconnectivity we observed in this study most likely reflects the hyperplastic effects of PSI to enhance synaptogenesis and dendritic growth. Does the concomitant increase in the blood level of small lipids reflect these morphological changes occurring in neuronal plasma membranes? Indeed, a majority of the brain lipidome composition in the prefrontal cortex of humans, chimpanzees, and macaque monkeys occur prior to adulthood, with significant alterations in lipid concentrations during early development^124^. Lipidomic studies in early childhood have shown marked changes in circulating lipid profiles from birth through early childhood, with specific lipid species showing age-dependent changes^125^. These findings collectively indicate that both postnatal and adolescent periods are characterized by enhanced lipidomic activity in plasma and brain, reflecting critical phases of neurodevelopment and metabolic regulation.

## Summary

Repetitive mild head injury affects diverse populations across age groups and is a significant risk factor for neurodegenerative diseases. Our rat model replicates clinical mild TBI by delivering impacts to awake animals during their active phase, producing no structural damage but causing measurable neuropathology. While we observed no significant behavioral deficits—consistent with rapid recovery in mild TBI—our model revealed several key pathological changes.

Diffusion imaging showed widespread increases in ADC values, suggestive of transient vasogenic edema. PSI treatment reduced edema, potentially through two mechanisms: 1) strengthening blood-brain barrier integrity and 2) enhancing astrocytic fluid clearance through GFAP-mediated changes in cell volume regulation.

CO2 challenge revealed lasting vascular hyperreactivity three weeks after injury, contrasting with the hyporeactivity seen in severe TBI. This suggests compensatory regulation in mild injury. PSI partially normalized this response, particularly in the basal ganglia, prefrontal cortex, and olfactory regions.

Functional connectivity analysis demonstrated global hypoconnectivity post-injury, matching clinical observations in head-injured patients. PSI treatment induced dramatic hyperconnectivity, notably in dopaminergic pathways to thalamic and somatosensory regions. This finding is particularly relevant given the role of rmTBI in risk for Parkinson’s disease.

Protein analysis showed an increase in BDNF and TrkB, a possible mechanism of PSI action. BDNF is crucial for neuronal survival and plasticity and acts through TrkB receptors, which increase after brain injury. Recent research shows PSI can directly bind TrkB with high affinity. The ability of PSI to reduce tau phosphorylation suggests potential therapeutic applications beyond rmTBI, possibly extending to other tau-related neurodegenerative disorders.

Modulations in plasma lipids with both rmTBI alone and in conjunction with PSI treatment provide a novel set of biomarkers to exploit for our understanding of how systemic changes in lipid signaling are associated with both long-term damage of rmTBI and the therapeutic effects of psilocybin.

This translational model successfully bridges bench-to-bedside by replicating clinical observations and identifies PSI as a promising therapeutic agent for repetitive mild head injury and its neurodegenerative consequences.

### Conflict of Interest

C.F.F. and P.P.K. have a partnership interest in Ekam Imaging Inc., a company that develops RF electronics and 3D MRI atlases for animal research. None of the other authors have a conflict of interest.

### Funding

Ekam Imaging Inc. The Paul H. Boerger Fund of the Delaware Community Foundation. Fellowships for M.I.A.: NIH-NIGMS R25GM122722 and National Science Foundation 2023004

### Authors’ contributions

All contributed equally to the data generation, while E.K.B., B.A., A.M., R.J.O., R.U., and P.P.K. were responsible for MRI and behavior data analysis. H.B.B. and T.J.W. performed blood analysis for lipidomics. M.A.G. and M.I.A. ran studies on molecular biology. E.K.B., P.P.K., H.B.B., M.A.G., and C.F.F. contributed to the concept, experimental design, manuscript preparation, and interpretation.

## Acknowledgements

We thank the National Institute on Drug Abuse Drug Supply Program and the Research Triangle Institute for providing the psilocybin used in these studies. We also thank the Center for Translational NeuroImaging and Ekam Imaging for financial support.

## Notes

### Summary of Updates

Figure 5 and corresponding results revised

